# Sources of Intraspecific Morphological Variation in *Vipera seoanei*: Allometry, Sex, and Colour Phenotype

**DOI:** 10.1101/2020.04.23.058206

**Authors:** Nahla Lucchini, Antigoni Kaliontzopoulou, Guillermo Aguado Val, Fernando Martínez-Freiría

## Abstract

Snakes frequently exhibit ontogenetic and sexual variation in head dimensions, as well as the occurrence of distinct colour morphotypes which might be fitness-related. In this study, we used linear biometry and geometric morphometrics to investigate intraspecific morphological variation related to allometry and sexual dimorphism in *Vipera seoanei*, a species that exhibits five colour morphotypes, potentially subjected to distinct ecological pressures. We measured body size (SVL), tail length and head dimensions in 391 specimens, and examined variation in biometric traits with respect to allometry, sex and colour morph. In addition, we analysed head shape variation by recording the position of 29 landmarks in 123 specimens and establishing a low-error protocol for implementing geometric morphometrics to European vipers. All head dimensions exhibited significant allometry, while sexual differences occurred for SVL, relative tail length and snout height. After considering size effects, we found significant differences in body proportions between the sexes and across colour morphs, which suggests an important influence of lowland and montane habitats in shaping morphological variation. By contrast, head shape did not exhibit significant variation across sexes or colour morphs. Instead it was mainly associated to allometric variation, where the supraocular and the rear regions of the head were the areas that varied the most throughout growth and across individuals. Overall, this study provides a thorough description of morphological variability in *Vipera seoanei* and highlights the relevance of combining different tools (i.e. linear and geometric morphometrics) and analyses to evaluate the relative contribution of different factors in shaping intraspecific variation.

## Introduction

Understanding how morphological variation across individuals arises and how it is distributed both temporally and spatially have attracted the attention of biologists over the years (Verwaijen, Van Damme and Herrel, 2002; Harmon et al., 2003; Kaliontzopoulou, Pinho and Martínez-Freiría, 2018).

Size is the predominant axis of morphological variation within and among populations (Rohlf, 1990). As such, allometry, the dependence of shape on size, is a major framework for understanding how and why different traits vary (Klingenberg, 2016). First, changes in size and shape occurring during growth and their relationship (i.e. ontogenetic allometry) are of essential importance for investigating the developmental processes producing the structures of interest (McNamara, 2012). Second, allometric variation across individuals at the same developmental stage within a population (i.e. static allometry) can be informative on the selective processes acting on individuals, since allometric parameters can be directly linked to both ecological adaptation (Gould, 1966) and sexual selection (Bonduriansky and Day, 2003). Indeed, body structures that are of particular relevance for either resource and/or mate acquisition, frequently tend to be positively allometric both ontogenetically and statically, a pattern which reflects higher investment during growth and a selective advantage for those individuals that possess a larger relative size of such structures, respectively.

Likewise, sexual dimorphism, is probably the second most important source of morphological variation in many animal species (Shine, 1989, 1994; Bonduriansky, 2007). Because of their different reproductive roles, adult males and females frequently exhibit different morphologies, which – from an evolutionary perspective – reflect the combined result of sexual and natural selection on members of each sex (Slatkin, 1984; Clutton-Brock, 2007). Under such influences, total body size, as well as the relative size of structures that enhance the reproductive potential of individuals, reach different evolutionary optima in members of each sex, depending on the social and/or ecological requirements of males and females (Kodric-Brown, Sibly, and Brown, 2006; Rodriguez et al., 2015). From a more developmental standing point, morphological differences between the sexes frequently arise as a result of highly divergent growth between males and females (Badyaev, 2002). As such, combining analyses that consider size and sexual variation simultaneously can be informative on the proximate and evolutionary causes that drive intraspecific morphological variation.

Due to their simple structure and organization, reptiles are excellent model organisms for morphological research (Gaffney, 1979; Forsman, 1996; Lovern, Holmes and Wade, 2004; Barata et al., 2012). Lack of limbs makes snakes even simpler than other reptiles and, thus, they are particularly popular subjects for studies of allometry (Shine, 1994; Feldman and Meiri, 2013). Snakes frequently show ontogenetic, sexual and phylogenetic variation in head dimensions relative to body size (Greene, 1983; Forsman, 1991; Shine, 1991; King, 1997). Similarly, snakes exhibit remarkable variation in the degree of sexual dimorphism and many studies have focused on sexual size and shape dimorphism and its evolutionary role in this group (Shine, 1994; Fairbairn, Blanckenhorn and Szekely, 2007; Krause, Burghardt and Gillingham, 2003; Henao-Duque and Ceballos, 2013). The most frequently evoked explanation for the evolution of sexual dimorphism is sexual selection acting through female choice or male-male interaction (Darwin, 1871). In the latter case (male-male competition) a larger body size (and head) in males is thought to be favoured by selection as it provides a competitive advantage over other individuals in combat or resource defence (Fairbairn, Blanckenhorn and Szekely, 2007). However, an alternative hypothesis, not necessarily exclusive of the first, considers natural selection acting to reduce competition between the sexes (Shine, 1991).

Colouration is another important type of intraspecific morphological variation in snakes, which is of great interest in different fields of study (Forsman, 1995a, 1995b; Zuffi, 2008; Martinez-Freiría et al., 2017). Colour pattern is an important component of a snake’s morphology, known to be fitness-related in many species, as it is crucial for fulfilling tasks necessary for their survival. Indeed, crypsis, Batesian mimicry or aposematism are ecological functions linked to the dorsal pattern of snakes, which determine how individuals interact with predators, prey and the surrounding environment (e.g. Valkonen et al., 2011; Santos et al., 2017; Martinez-Freiría et al., 2017). Similarly, dark colouration plays an important role in thermoregulation in ectotherms inhabiting cold environments (i.e. *thermal melanism hypothesis;* Clusella-Trullas et al., 2008). Accordingly, melanistic snakes can exhibit increased growth rates, better body condition and locomotor performance, longer activity periods, or higher fertility in females than non-melanistic ones (Luiselli, 1992; Capula and Luiselli, 1994; Castella et al., 2013).

The Iberian adder, *Vipera seoanei* (Lataste, 1879), is a small-sized venomous snake with Euro-Siberian affinity, nearly endemic to the Iberian Peninsula (Martínez-Freiría and Brito, 2014). It is distributed across north-western Portugal and northern Spain, with some populations penetrating a few kilometres into south-western France (Braña, 2002; Brito, 2008; Martínez-Freiría and Brito, 2014). *Vipera seoanei* exhibits low intraspecific phylogenetic variability and is characterized by high levels of polymorphism in pholidotic traits and colour morphs (Bea et al., 1984; Saint-Girons, Braña and Bea, 1986; Martínez-Freiría and Brito, 2013; Martínez-Freiría, Velo-Antón and Brito, 2015, Martínez-Freiría et al., 2017). Pholidotic trait variability correlates to variation in climatic conditions across the species’ range and, thus, these traits were suggested to be under selection or to be shaped by plasticity (Martínez-Freiría and Brito, 2013).

Remarkably, *V. seoanei* exhibits a pronounced level of polymorphism, with five colour morphs which, in general terms, differ in their geographical distributions (see Martínez-Freiría et al., 2017): *bilineata* (brownish or reddish, with a wide dark dorsal line) located in the central coastal and mountain Cantabrian region; *classic* (brownish or reddish, with a wide dorsal zig-zag pattern) found all over the lowland species range; *cantabrica* (greyish, with a dorsal line and a triangular zig-zag or transversal bars pattern) mostly restricted to the Cantabrian Mountains; *uniform* (greyish or light brown, with no dorsal pattern) restricted to the eastern Cantabrian Mountains; and *melanistic* (completely black, with no dorsal pattern) generally occurring at high elevation ranges. Predation and thermal pressures were found to be related to the frequency of these morphs within populations (Martínez-Freiría et al., 2017). However, colouration types span across contrasting environments (in terms of climate, prey and predators) which are likely to affect their morphological properties. Specifically, lowland habitats, characterized by a milder climate than montane ones, could allow vipers to present an extended cycle of activity, with more prey intake and higher yearly growth rates (e.g. in *Vipera aspis*; Zuffi et al., 2009). Consequently, earlier sexual maturity and larger sizes could be expected for individuals inhabiting lowlands. Conversely, predation pressures are suggested to be lower in montane than in lowland regions (see Martínez-Freiría et al., 2017), and, thus, survival of individuals in montane regions is expected to be higher, allowing vipers to reach larger sizes compared to individuals in the lowlands.

In this work, we combine linear biometry and geometric morphometrics to describe intraspecific morphological variation in *Vipera seoanei* and provide an integrated assessment of the relevance of allometry, sexual dimorphism and colour phenotype in shaping body size, body shape, and head shape variation. Specifically, we set out to investigate the following questions: 1) Is there morphological sexual dimorphism within *V. seoanei*? 2) Do different colour morphs exhibit different morphological properties? and 3) What is the role of allometric variation in moulding morphological variation among individuals of different sexes and colour morphs? By addressing these questions, we aim at providing a detailed evaluation of the relative importance of these major sources of variation in determining morphological variability within this emblematic viper.

## Materials and Methods

### Specimens

We analysed a total of 391 specimens of *V. seoanei*, coming from the whole distributional range of the species (Fig. 1), which resulted from a careful selection from over 500 specimens, where any damaged specimens were excluded. Of these analysed specimens, 61 were collected in the field and measured alive, 35 were found road killed in the field, and 295 were specimens preserved in eight Natural History Museum collections: Estación Biológica de Doñana - CSIC (Sevilla, Spain), Museo Nacional de Ciencias Naturales -CSIC (Madrid, Spain), Sociedad de Ciencias Aranzandi (Donostia, Spain), Universidad de Oviedo (Oviedo, Spain), Universidad de Salamanca (Salamanca, Spain), Universidade da Coruña (A Coruña, Spain), Universidade de Santiago de Compostela -Museo Luis Iglesias (Santiago de Compostela, Spain) and Zoologisches Forschungsmuseum Alexander Koenig (Bonn, Germany). Vipers were divided into six sex/age groups (adult males, adult females, subadult males, subadult females, juvenile males and juvenile females) used exclusively in order to substitute the missing values of each linear biometric trait with the corresponding group mean (see Table S1.3 in Supp. Mat. S1). Age groups were based on size categories divided as follow: juveniles SVL< 250 mm, subadults 250 mm < SVL > 325 mm, adults SVL > 325 mm. Information on growth and sexual maturity for *V. seoanei* is very limited (see Martínez-Freiría and Brito, 2014) and thus, size categories were based on information available for other Iberian vipers (e.g. Martínez-Freiría et al., 2010), as well as, our experience with the species. Age groups, however, were not further considered, as size-related variation was represented through body size (SVL) for subsequent analyses. Additionally, we classified each specimen in one of five colour morphs (bilineata, classic, cantabrica, melanistic and uniform) following criteria used in previous works (Martínez-Freiría and Brito, 2013; Martínez-Freiría et al., 2017). The uniform type was very rare in our dataset (n = 6; Supp. Mat. 1). Therefore, in the analyses of colour morphs, individuals showing this pattern were removed and this pattern type was not considered.

**Figure 1.**
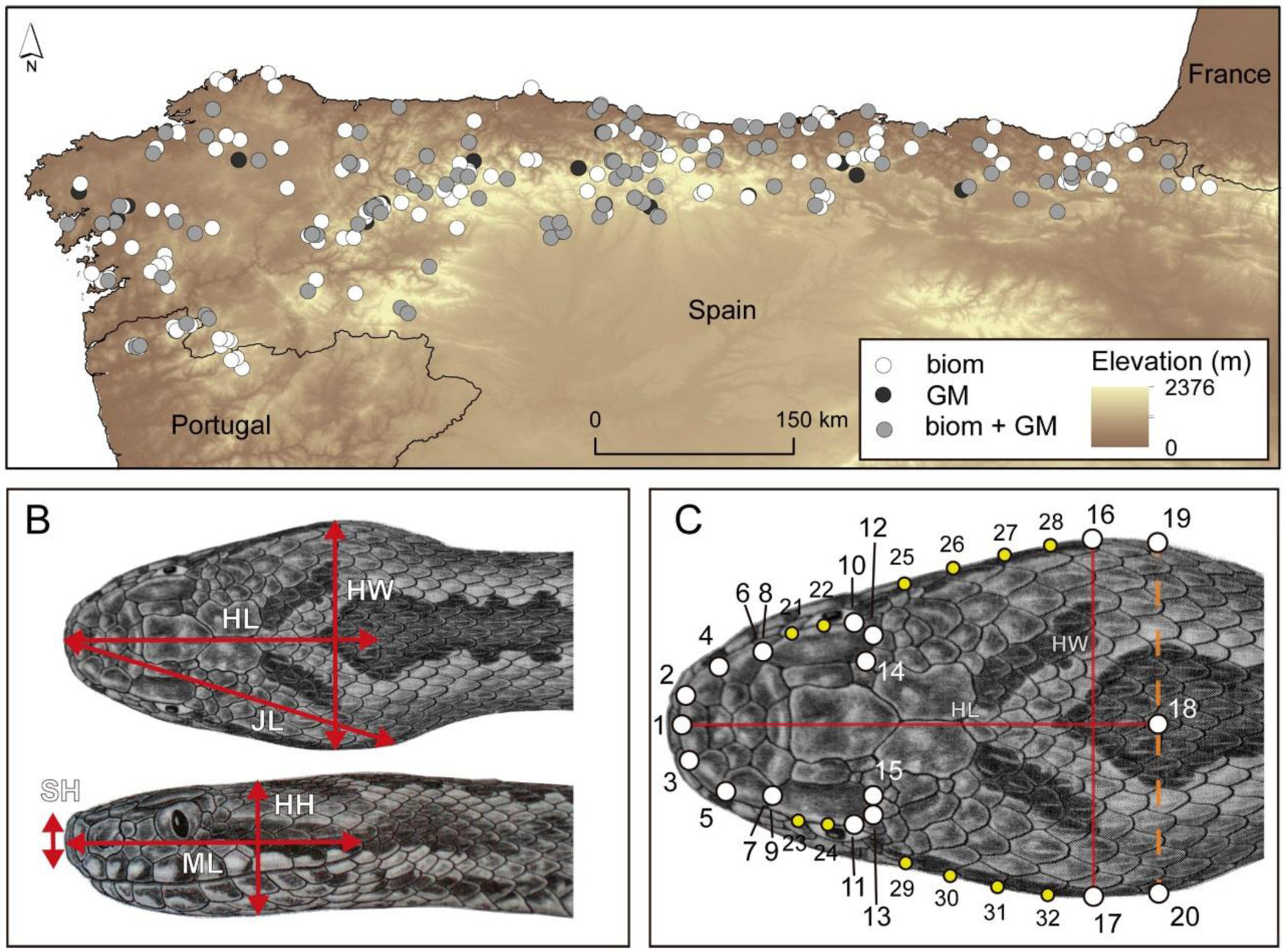
(A) distribution of specimens of *V. seoanei* considered in this study, including specimens used for biometric (white-filled dots) or GM (black-filled dots) analyses only, and specimens used for both analyses (grey-filled dots); (B) linear measurements in the dorsal and lateral views of the head (HL, head length; HW, head width; JL, jaw length; SH, snout height; HH, head height; ML, mouth length); (C) position of landmarks (white-filled dots) and semi-landmarks (yellow-filled dots) in the dorsal view of the head. Head length (HL) and width (HW) are also depicted as they were used to calibrate images (see Suppl. Mat. S1 for details on landmarks and semi-landmarks). Drawings of the head authored by Pedro Salgado (from Fauna Ibérica vol. 24, MNCN-CSIC).

In order to analyse morphological variation among individuals of this species, we used two different approaches: linear biometry, applied to the whole body of the analysed individuals, and geometric morphometrics, applied to quantify head shape. For the first approach, a total of 391 specimens (191 females and 200 males) were considered, while only 124 specimens (58 females and 66 males) were considered for geometric morphometrics (see Table S1.1 and S1.2 in Supp. Mat. S1 for details).

### Linear biometry

For each specimen, we considered the following biometric traits (Fig. 1): snout-vent length (SVL), tail length (TAIL), head length (HL), mouth length (ML), jaw length (JL), head width (HW), head height (HH), and snout height (SH), measured to the closest 0.01mm using electronic callipers or a ruler. All measurements were log-transformed prior to analyses.

Based on these variables, we first performed an Analysis of Variance (ANOVA) considering all specimens to test for differences between the sexes in morphometric measures in an ontogenetic context. For this purpose, we used sex as a fixed factor and SVL as a covariate, also considering the interaction between sex and SVL to test for differences in allometric slopes between the sexes.

Next, we performed an ANOVA where we examined variation in biometric traits with respect to sex and colour morphs among adults. For this purpose, we first performed an ANOVA to analyse variation in body size (SVL) across sexes and colour morphs. Then, we performed ANCOVA comparisons as described for the ontogenetic analyses to investigate static allometric variation in the rest of the morphometric characters, using sex and colour as fixed effects, SVL as the covariate, and including also all interaction terms.

### Geometric morphometrics

To quantify variation in head shape and size, a two-dimensions geometric morphometric approach was applied on photos of the dorsal view of the head.

Twenty-one fixed landmarks and eight semi-landmarks (Gunz et al., 2009) were digitalized for each specimen using the program tpsDig2 version 2.29 (Rohlf, 2008), resulting in 29 pairs of coordinates (Fig. 1; Table 1). This process was repeated twice to quantify digitizing error induced by the observer and verify that the origin of the specimens did not influence digitizing procedures (see Suppl. Mat. S2). When the position of the landmarks could not be precisely defined, their coordinates were considered as missing data and their values predicted through a linear regression model (Gunz et al., 2009; Arbour and Brown, 2014). After data acquisition, a Generalized Procrustes Analysis (GPA, Rolhf and Slice, 1990; Rolhf, 1999) was applied in order to standardize size, location and rotation effects, and to preserve only information related to shape (Dryden and Mardia, 1998; Claude, 2008). During this process the position of each semi-landmark was optimized by sliding it along its tangent line, using the criterion of minimizing the Procrustes distance between specimens (Bookstein, 1997), which does not influence the position of the rest of landmarks and semi-landmarks (Gunz and Mitteroecker, 2013). Since we were not interested in evaluating asymmetry, we only considered the symmetry shape component for each individual, calculated as the mean of the coordinates of the two sides of the head across the midline (Klingenberg, Barluenga and Meyer, 2002). To take digitizing error into account, we used the average of the two repetitions.

**Table 1:** List and description of landmarks and semi-landmarks used for quantifying head shape in *V. seoanei* (see Figure 1C).

|  |  | Description |
| --- | --- | --- |
| **Landmark** | 1 | Anterior point of the joint between the two apical scales. |
|  | 2, 3 | External margin of the joint between the apicals and the first canthal scale. |
|  | 4, 5 | External margin of the joint between the first and second canthal. |
|  | 6, 7 | External posterior border of the second canthal. |
|  | 8, 9 | External anterior corner of the supraocular. |
|  | 10, 11 | External point of contact between the supraocular and the periocular. |
|  | 12, 13 | Middle of the posterior border of the supraocular, where it contacts with the posterior border of the periocular. |
|  | 14, 15 | Internal posterior border of the supraocular. |
|  | 16, 17 | External margins of the widest region of the head*. |
|  | 18 | On the vector perpendicular to the line of landmarks # 16-17, at distance equal to head length (HL) from the snout. |
|  | 19, 20 | On the border of the head, at the line perpendicular to landmarks # 1-10, and passing through landmark # 10. |
| **Semilandmark** | 21 - 24 | Semilandmarks between landmarks # 8-10 and 9-11, correspondingly, defining the outline of the supraocular scale. |
|  | 25 - 32 | Semilandmarks between landmarks # 10-16 and 11-17, correspondingly, defining the outline of the posterior region of the head. |

\*Images were calibrated using the distance between landmarks # 16 and 17, which corresponds to head width (HW)

As for linear biometric variables, ontogenetic and static allometry were explored through Procrustes ANOVAs with permutation procedures (1000 permutations). First, we used the entire dataset to explore ontogenetic variation and examine its interaction with sex and pattern type through an ANOVA that included sex and pattern as fixed factors, and the logarithm of centroid size (logCS) as the covariate, as well as all interaction terms. Second, we focused on static allometric shape variation only in adults. Here, we first performed ANOVA comparisons considering sex, pattern type and their interaction to test for differences among groups in head size (logCS); then, we examined variation in head shape with respect to head size across groups by fitting a linear model with sex and pattern type as fixed effects and logCS as the covariate.

To further explore and describe variation in head shape across specimens, we performed a principal component analysis (PCA) of shape variables and visualized the associated shape variation using deformation grids to depict shape change across the extremes of PC axes. Then, head shape was size-corrected through a linear model based on logCS, and a second PCA was performed and deformation grids produced to visualize shape variation after removing size effects.

All analyses were performed using the R language for statistical programming (R Core Team, 2011), and all geometric morphometrics procedures were implemented in the *geomorph* R-package (Adams, Collyer and Kaliontzopoulou, 2019). All ANOVA procedures were evaluated for significance using residual-randomization permutation procedures as implemented in the R-package *RRPP* (Collyer and Adams 2018, 2019).

## Results

### Linear biometry

Descriptive statistics for the individuals of males and females of *Vipera seoanei* examined here, for each biometric trait considered can be found in Supplementary Table S1.4. In ontogenetic allometry analyses, the linear models examined indicated that in *Vipera seoanei* SVL significantly influenced all the biometric characteristics considered (Table 2). However, we did not find a significant influence of sex on most biometric variables. Significant differences between the sexes were observed only for snout height, and when considering sex and the interaction between sex and SVL for tail length (Table 2, Fig. 2). When focusing only on adult individuals and including also colour morph, we found a significant effect of sex, pattern and their interaction on body size (SVL) (Table 3). Specifically, pairwise comparisons based on 1000 permutations of Euclidean distances indicated that differences across colour morphs within each sex were only significant between *cantabrica* and *classic* for females, where *cantabrica* exhibited larger body size; and between *bilineata* and *classic* in males, where *bilineata* exhibited smaller body size (Fig. 2; Supp. Mat. 3).

**Table 2.** Results of ANOVA considering all specimens at the ontogenetic level. df: degrees of freedom, SS: Sums of Squares, MS: Mean Squares, R^2^: coefficient of determination, F: F-value, Z: standardized z-score, p: corresponding p-value based on 1000 permutations of residuals. TAIL (tail length), HL (head length), ML (mouth length), JL (jaw length), HW (head width), HH (head height), SH (snout height). Significant p-values are marked in bold.

|  |  | df | SS | MS | R<sup>2</sup> | F | Z | p |
| --- | --- | --- | --- | --- | --- | --- | --- | --- |
| TAIL | svl | 1 | 21.487 | 21.487 | 0.766 | 2940.372 | 4.142 | **0.001** |
|  | sex | 1 | 3.755 | 3.755 | 0.134 | 513.883 | 3.347 | **0.001** |
|  | svl×sex | 1 | 0.075 | 0.075 | 0.003 | 10.329 | 1.590 | **0.003** |
|  | Residuals | 376 | 2.748 | 0.007 | 0.098 |  |  |  |
|  | Total | 379 | 28.065 |  |  |  |  |  |
| HL | svl | 1 | 8.057 | 8.057 | 0.836 | 1941.552 | 4.039 | **0.001** |
|  | sex | 1 | 0.000 | 0.000 | 0.000 | 0.009 | -1.633 | 0.925 |
|  | svl×sex | 1 | 0.000 | 0.000 | 0.000 | 0.022 | -1.162 | 0.876 |
|  | Residuals | 382 | 1.585 | 0.004 | 0.164 |  |  |  |
|  | Total | 385 | 9.643 |  |  |  |  |  |
| ML | svl | 1 | 8.661 | 8.661 | 0.921 | 4351.974 | 4.296 | **0.001** |
|  | sex | 1 | 0.004 | 0.004 | 0.000 | 1.840 | 0.836 | 0.192 |
|  | svl×sex | 1 | 0.006 | 0.006 | 0.001 | 3.208 | 1.132 | 0.074 |
|  | Residuals | 367 | 0.730 | 0.002 | 0.078 |  |  |  |
|  | Total | 370 | 9.401 |  |  |  |  |  |
| JL | svl | 1 | 9.114 | 9.114 | 0.914 | 3868.491 | 4.185 | **0.001** |
|  | sex | 1 | 0.000 | 0.000 | 0.000 | 0.009 | -1.376 | 0.914 |
|  | svl×sex | 1 | 0.007 | 0.007 | 0.001 | 3.175 | 1.154 | 0.082 |
|  | Residuals | 363 | 0.855 | 0.002 | 0.086 |  |  |  |
|  | Total | 366 | 9.977 |  |  |  |  |  |
| HW | svl | 1 | 9.338 | 9.338 | 0.737 | 937.380 | 3.819 | **0.001** |
|  | sex | 1 | 0.000 | 0.000 | 0.000 | 0.001 | -2.666 | 0.984 |
|  | svl×sex | 1 | 0.017 | 0.017 | 0.001 | 1.732 | 0.817 | 0.195 |
|  | Residuals | 332 | 3.307 | 0.010 | 0.261 |  |  |  |
|  | Total | 335 | 12.663 |  |  |  |  |  |
| HH | svl | 1 | 8.988 | 8.988 | 0.790 | 1191.485 | 3.662 | **0.001** |
|  | sex | 1 | 0.008 | 0.008 | 0.001 | 1.036 | 0.606 | 0.298 |
|  | svl×sex | 1 | 0.017 | 0.017 | 0.001 | 2.243 | 0.906 | 0.149 |
|  | Residuals | 314 | 2.369 | 0.008 | 0.208 |  |  |  |
|  | Total | 317 | 11.381 |  |  |  |  |  |
| SH | svl | 1 | 7.954 | 7.954 | 0.784 | 1249.119 | 3.651 | **0.001** |
|  | sex | 1 | 0.050 | 0.050 | 0.005 | 7.850 | 1.518 | **0.006** |
|  | svl×sex | 1 | 0.001 | 0.001 | 0.000 | 0.143 | -0.310 | 0.710 |
|  | Residuals | 337 | 2.146 | 0.006 | 0.211 |  |  |  |
|  | Total | 340 | 10.150 |  |  |  |  |  |

**Table 3.**
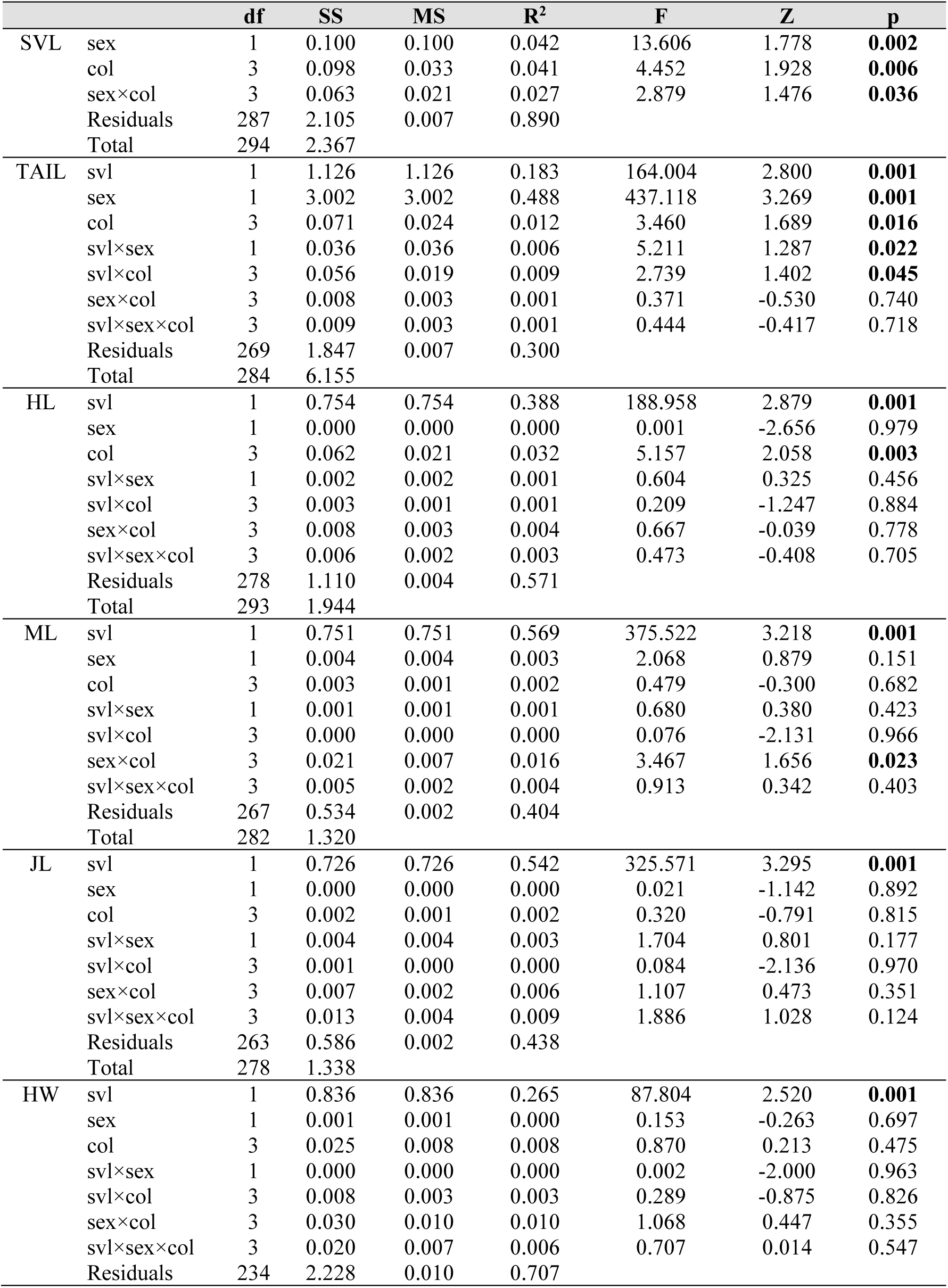
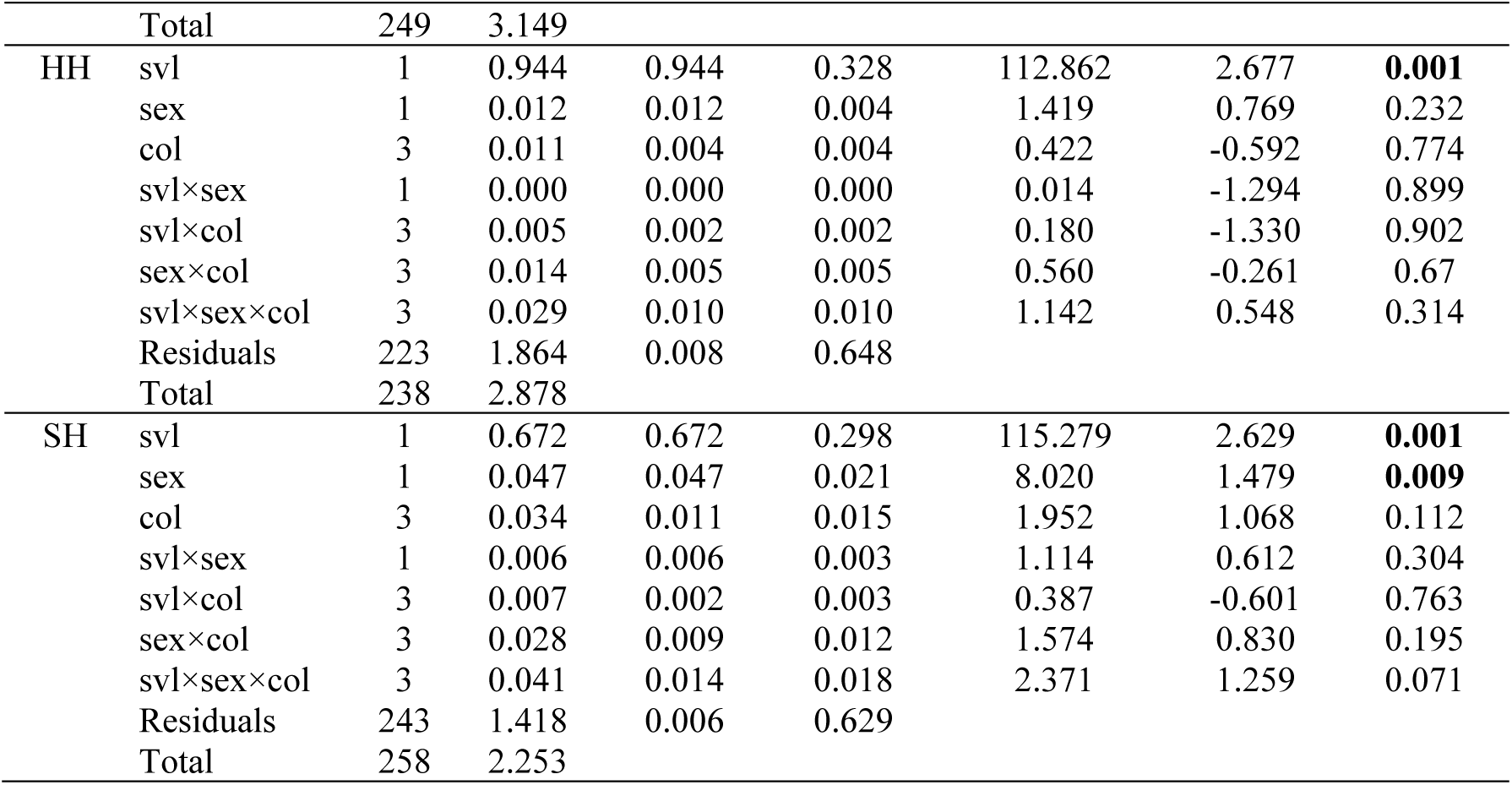
Results of ANOVA considering only adults to examine variation related to size, sex and colour morph (col). df: degrees of freedom, SS: Sums of Squares, MS: Mean Squares, R^2^: coefficient of determination, F: F-value, Z: standardized z-score, p: corresponding p-value based on 1000 permutations of residuals. SVL (snout-vent length), TAIL (tail length), HL (head length), ML (mouth length), JL (jaw length), HW (head width), HH (head height), SH (snout height). Significant p-values are marked in bold.

|  |  | df | SS | MS | R<sup>2</sup> | F | Z | p |
| --- | --- | --- | --- | --- | --- | --- | --- | --- |
| SVL | sex | 1 | 0.100 | 0.100 | 0.042 | 13.606 | 1.778 | **0.002** |
|  | col | 3 | 0.098 | 0.033 | 0.041 | 4.452 | 1.928 | **0.006** |
|  | sex×col | 3 | 0.063 | 0.021 | 0.027 | 2.879 | 1.476 | **0.036** |
|  | Residuals | 287 | 2.105 | 0.007 | 0.890 |  |  |  |
|  | Total | 294 | 2.367 |  |  |  |  |  |
| TAIL | svl | 1 | 1.126 | 1.126 | 0.183 | 164.004 | 2.800 | **0.001** |
|  | sex | 1 | 3.002 | 3.002 | 0.488 | 437.118 | 3.269 | **0.001** |
|  | col | 3 | 0.071 | 0.024 | 0.012 | 3.460 | 1.689 | **0.016** |
|  | svl×sex | 1 | 0.036 | 0.036 | 0.006 | 5.211 | 1.287 | **0.022** |
|  | svl×col | 3 | 0.056 | 0.019 | 0.009 | 2.739 | 1.402 | **0.045** |
|  | sex×col | 3 | 0.008 | 0.003 | 0.001 | 0.371 | -0.530 | 0.740 |
|  | svl×sex×col | 3 | 0.009 | 0.003 | 0.001 | 0.444 | -0.417 | 0.718 |
|  | Residuals | 269 | 1.847 | 0.007 | 0.300 |  |  |  |
|  | Total | 284 | 6.155 |  |  |  |  |  |
| HL | svl | 1 | 0.754 | 0.754 | 0.388 | 188.958 | 2.879 | **0.001** |
|  | sex | 1 | 0.000 | 0.000 | 0.000 | 0.001 | -2.656 | 0.979 |
|  | col | 3 | 0.062 | 0.021 | 0.032 | 5.157 | 2.058 | **0.003** |
|  | svl×sex | 1 | 0.002 | 0.002 | 0.001 | 0.604 | 0.325 | 0.456 |
|  | svl×col | 3 | 0.003 | 0.001 | 0.001 | 0.209 | -1.247 | 0.884 |
|  | sex×col | 3 | 0.008 | 0.003 | 0.004 | 0.667 | -0.039 | 0.778 |
|  | svl×sex×col | 3 | 0.006 | 0.002 | 0.003 | 0.473 | -0.408 | 0.705 |
|  | Residuals | 278 | 1.110 | 0.004 | 0.571 |  |  |  |
|  | Total | 293 | 1.944 |  |  |  |  |  |
| ML | svl | 1 | 0.751 | 0.751 | 0.569 | 375.522 | 3.218 | **0.001** |
|  | sex | 1 | 0.004 | 0.004 | 0.003 | 2.068 | 0.879 | 0.151 |
|  | col | 3 | 0.003 | 0.001 | 0.002 | 0.479 | -0.300 | 0.682 |
|  | svl×sex | 1 | 0.001 | 0.001 | 0.001 | 0.680 | 0.380 | 0.423 |
|  | svl×col | 3 | 0.000 | 0.000 | 0.000 | 0.076 | -2.131 | 0.966 |
|  | sex×col | 3 | 0.021 | 0.007 | 0.016 | 3.467 | 1.656 | **0.023** |
|  | svl×sex×col | 3 | 0.005 | 0.002 | 0.004 | 0.913 | 0.342 | 0.403 |
|  | Residuals | 267 | 0.534 | 0.002 | 0.404 |  |  |  |
|  | Total | 282 | 1.320 |  |  |  |  |  |
| JL | svl | 1 | 0.726 | 0.726 | 0.542 | 325.571 | 3.295 | **0.001** |
|  | sex | 1 | 0.000 | 0.000 | 0.000 | 0.021 | -1.142 | 0.892 |
|  | col | 3 | 0.002 | 0.001 | 0.002 | 0.320 | -0.791 | 0.815 |
|  | svl×sex | 1 | 0.004 | 0.004 | 0.003 | 1.704 | 0.801 | 0.177 |
|  | svl×col | 3 | 0.001 | 0.000 | 0.000 | 0.084 | -2.136 | 0.970 |
|  | sex×col | 3 | 0.007 | 0.002 | 0.006 | 1.107 | 0.473 | 0.351 |
|  | svl×sex×col | 3 | 0.013 | 0.004 | 0.009 | 1.886 | 1.028 | 0.124 |
|  | Residuals | 263 | 0.586 | 0.002 | 0.438 |  |  |  |
|  | Total | 278 | 1.338 |  |  |  |  |  |
| HW | svl | 1 | 0.836 | 0.836 | 0.265 | 87.804 | 2.520 | **0.001** |
|  | sex | 1 | 0.001 | 0.001 | 0.000 | 0.153 | -0.263 | 0.697 |
|  | col | 3 | 0.025 | 0.008 | 0.008 | 0.870 | 0.213 | 0.475 |
|  | svl×sex | 1 | 0.000 | 0.000 | 0.000 | 0.002 | -2.000 | 0.963 |
|  | svl×col | 3 | 0.008 | 0.003 | 0.003 | 0.289 | -0.875 | 0.826 |
|  | sex×col | 3 | 0.030 | 0.010 | 0.010 | 1.068 | 0.447 | 0.355 |
|  | svl×sex×col | 3 | 0.020 | 0.007 | 0.006 | 0.707 | 0.014 | 0.547 |
|  | Residuals | 234 | 2.228 | 0.010 | 0.707 |  |  |  |
|  | Total | 249 | 3.149 |  |  |  |  |  |
| HH | svl | 1 | 0.944 | 0.944 | 0.328 | 112.862 | 2.677 | **0.001** |
|  | sex | 1 | 0.012 | 0.012 | 0.004 | 1.419 | 0.769 | 0.232 |
|  | col | 3 | 0.011 | 0.004 | 0.004 | 0.422 | -0.592 | 0.774 |
|  | svl×sex | 1 | 0.000 | 0.000 | 0.000 | 0.014 | -1.294 | 0.899 |
|  | svl×col | 3 | 0.005 | 0.002 | 0.002 | 0.180 | -1.330 | 0.902 |
|  | sex×col | 3 | 0.014 | 0.005 | 0.005 | 0.560 | -0.261 | 0.67 |
|  | svl×sex×col | 3 | 0.029 | 0.010 | 0.010 | 1.142 | 0.548 | 0.314 |
|  | Residuals | 223 | 1.864 | 0.008 | 0.648 |  |  |  |
|  | Total | 238 | 2.878 |  |  |  |  |  |
| SH | svl | 1 | 0.672 | 0.672 | 0.298 | 115.279 | 2.629 | **0.001** |
|  | sex | 1 | 0.047 | 0.047 | 0.021 | 8.020 | 1.479 | **0.009** |
|  | col | 3 | 0.034 | 0.011 | 0.015 | 1.952 | 1.068 | 0.112 |
|  | svl×sex | 1 | 0.006 | 0.006 | 0.003 | 1.114 | 0.612 | 0.304 |
|  | svl×col | 3 | 0.007 | 0.002 | 0.003 | 0.387 | -0.601 | 0.763 |
|  | sex×col | 3 | 0.028 | 0.009 | 0.012 | 1.574 | 0.830 | 0.195 |
|  | svl×sex×col | 3 | 0.041 | 0.014 | 0.018 | 2.371 | 1.259 | 0.071 |
|  | Residuals | 243 | 1.418 | 0.006 | 0.629 |  |  |  |
|  | Total | 258 | 2.253 |  |  |  |  |  |

**Figure 2.**
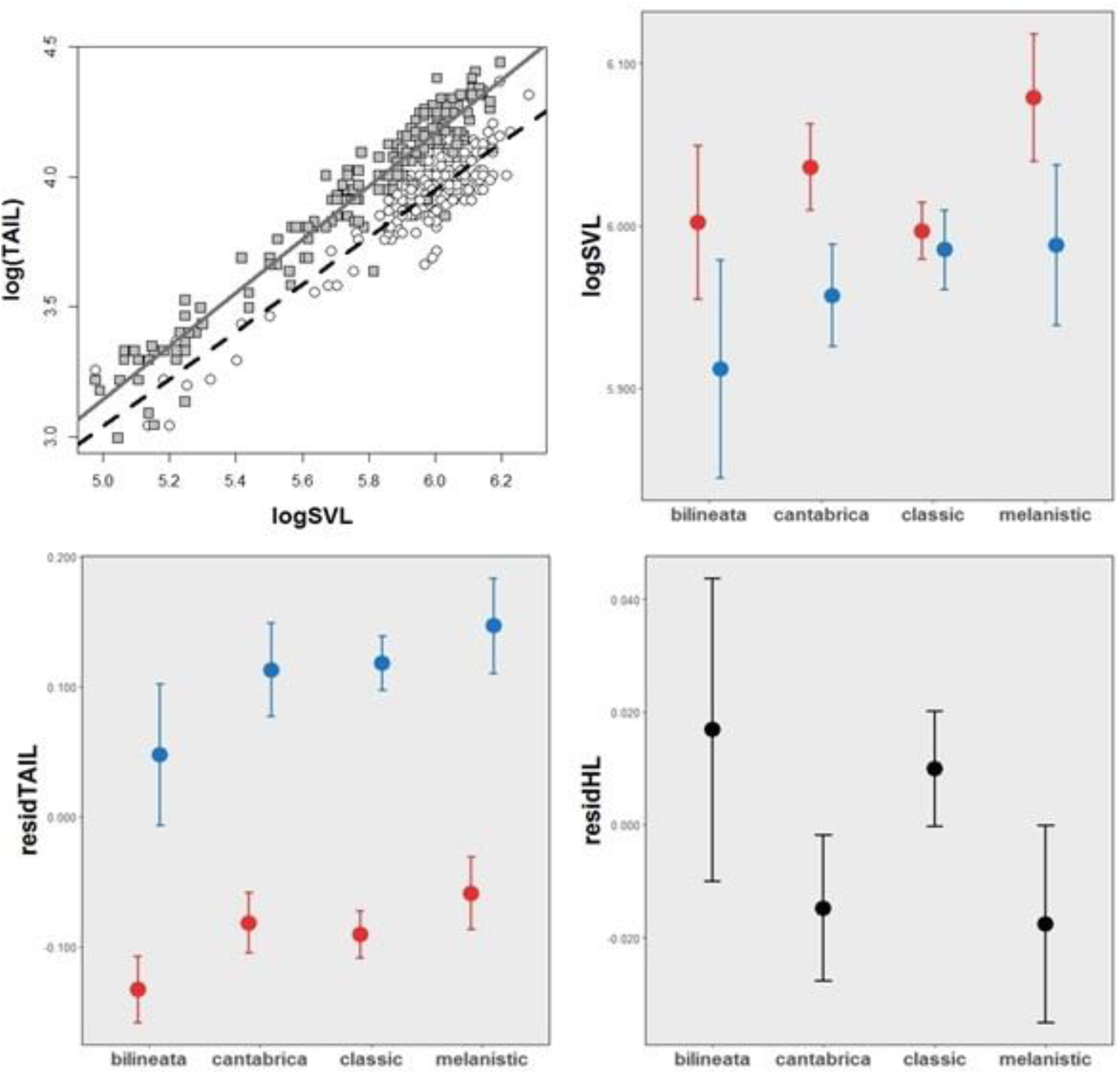
Ontogenetic relationship between tail length and body size (SVL) (top, left) and variation across colour morphs and sexes (red females, blue males) in biometric traits of *Vipera seoanei* (top, right, bottom left, bottom right). Grey squares, continuous line: males; white circles, dashed line: females. Points represent means and vertical bars denote 95% confidence intervals. LogSVL snout-vent length logarithm, residTAIL residual of tail length, residHL residual of head length. For residHL both sexes are pooled, as ANOVA comparisons did not reveal the existence of sexual dimorphism (see Table 2).

SVL had again a significant influence on all morphological measures (Table 3). Concerning static allometry analyses, we found significant effects of sex and colour morph on tail length and the interaction between SVL and sex and SVL and colour morph, as well as of colour morph – but not sex – on head length (Table 3). Considering relative tail length, pairwise comparisons based on 1000 permutations of Euclidean distances identified significantly longer tails in males of *classic* but not of other morphs (Fig. 2; Supp. Mat. 3). A similar pattern was identified for relative mouth length, for which *cantabrica* and *melanistic* females exhibited longer mouth than males, while head length didn’t give any significant results (Fig. 3; Supp. Mat. 3).

**Figure 3.**
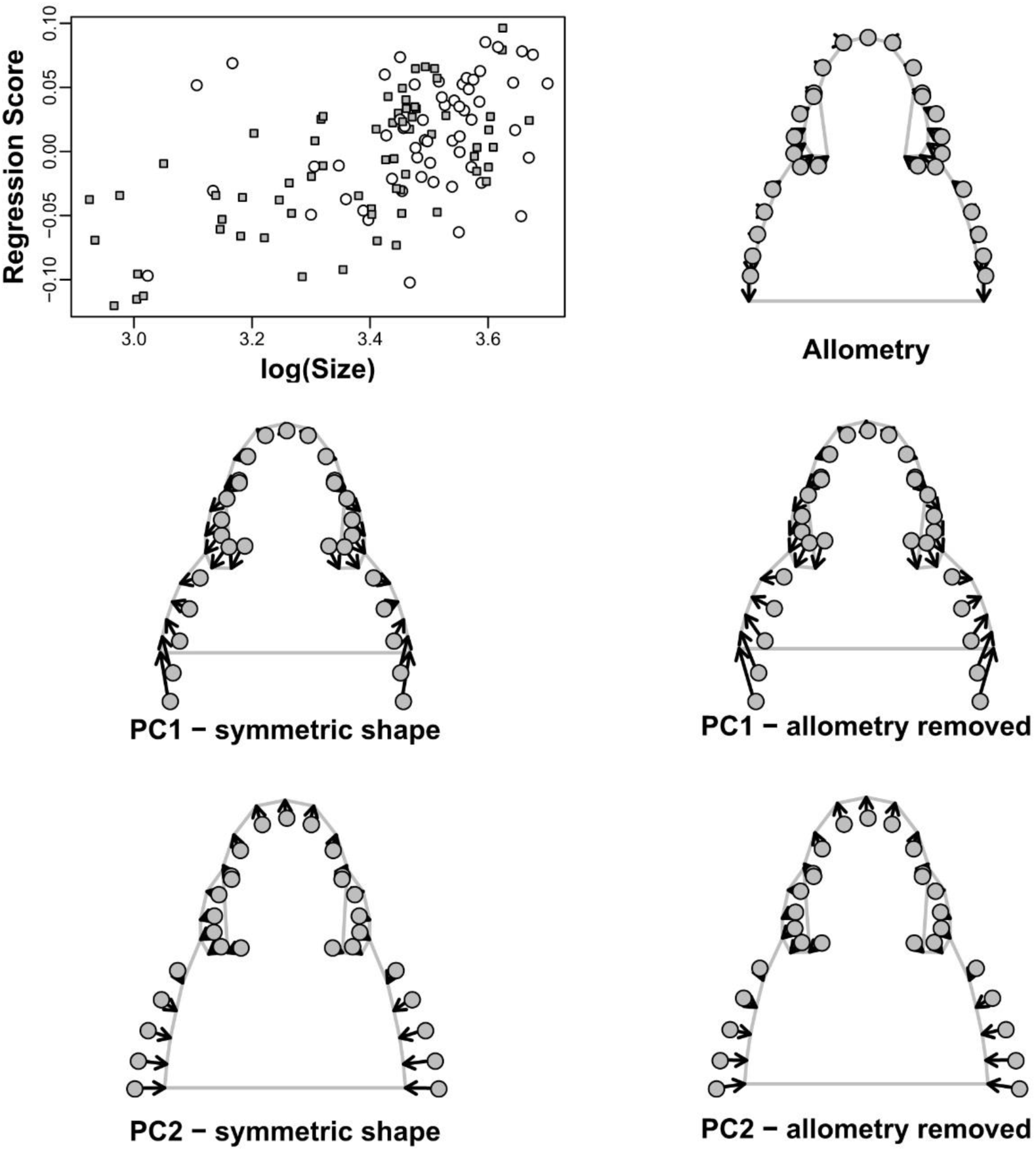
On the top right regression of shape variables on log centroid size, grey squares: males; white circles: females. On top left ontogenetic shape head variation. Shape change is magnified x2 to facilitate visualization. Center and bottom variation in head shape of *V. seoanei* across PC1 (center) and PC2 (bottom), before (left) and after (right) size correction, the length and direction of the vectors represent the change in shape from the negative to the positive extreme of each axis.

### Geometric morphometrics

The exploration of the symmetric component of head shape in the sample size revealed no significant differences and an absence of sexual dimorphism in *V. seoanei* (Table 4). When examining ontogenetic, we found a significant contribution of logCS in explaining variation in head shape and a contribution of the colour morph, while no significant effects were found for sex, colour morph or their interaction. In the static allometry only the logCS was significant (Table 4). Allometric variation in head shape was associated to a reduction of relative eye size and an amplification of the posterior region of the head throughout growth (Fig. 3), which occurred under the same pattern for both sexes.

**Table 4.** Results of ontogenetic and static allometry analyses of head shape. cs: centroid size. df: degrees of freedom, SS: Sums of Squares, MS: Mean Squares, R^2^: coefficient of determination, F: F-value, Z: standardized z-score, p: corresponding p-value based on 1000 permutations of residuals, col: colour morph. Significant p-values are marked in bold.

|  |  | df | SS | MS | R<sup>2</sup> | F | Z | p |
| --- | --- | --- | --- | --- | --- | --- | --- | --- |
| ONTOGENETIC | cs | 1 | 0.097 | 0.097 | 0.184 | 27.119 | 4.459 | **0.001** |
|  | sex | 1 | 0.004 | 0.004 | 0.007 | 1.080 | 0.488 | 0.332 |
|  | col | 3 | 0.020 | 0.007 | 0.039 | 1.902 | 1.724 | **0.046** |
|  | cs×sex | 1 | 0.001 | 0.001 | 0.003 | 0.374 | -0.844 | 0.793 |
|  | cs×col | 3 | 0.009 | 0.003 | 0.018 | 0.876 | -0.122 | 0.55 |
|  | sex×col | 3 | 0.006 | 0.002 | 0.012 | 0.568 | -0.887 | 0.805 |
|  | cs×sex×col | 3 | 0.007 | 0.002 | 0.012 | 0.612 | -0.857 | 0.796 |
|  | Residuals | 107 | 0.383 | 0.004 | 0.726 |  |  |  |
|  | Total | 122 | 0.528 |  |  |  |  |  |
| STATIC | cs | 1 | 0.029 | 0.029 | 0.086 | 8.515 | 3.046 | **0.001** |
|  | sex | 1 | 0.004 | 0.004 | 0.011 | 1.039 | 0.429 | 0.339 |
|  | col | 3 | 0.018 | 0.006 | 0.053 | 1.753 | 1.409 | 0.086 |
|  | cs×sex | 1 | 0.002 | 0.002 | 0.005 | 0.516 | -0.524 | 0.694 |
|  | cs×col | 3 | 0.009 | 0.003 | 0.026 | 0.840 | -0.117 | 0.568 |
|  | sex×col | 3 | 0.006 | 0.002 | 0.017 | 0.567 | -0.905 | 0.813 |
|  | cs×sex×col | 3 | 0.011 | 0.004 | 0.033 | 1.086 | 0.408 | 0.35 |
|  | Residuals | 76 | 0.259 | 0.003 | 0.769 |  |  |  |
|  | Total | 91 | 0.337 |  |  |  |  |  |

Principal Component Analysis (PCA) of shape variables retrieved a first component that explained 59.63% of the variability and captured shape variation mainly in the posterior head area and the relative size of the supraocular region, where an amplification of relative eye size as accompanied by a widening and shortening of the back of the head (Fig. 3). The second principal component summed up 20.26% of the variability, and it was associated to a concomitant amplification of the snout and the eyes, together with a reduction of relative head width (Fig. 3). Once shape variables were corrected for allometric effects, the variation captured by PC axes was slightly reduced (53.68% for PC1 and 23.54% for PC2), but global shape variation patterns remained unaltered (Fig. 3, centre and bottom, right).

## Discussion

In this study we investigated intraspecific morphological variability in *Vipera seoanei*, aiming to provide an assessment of how different factors mould body and head size and shape. We applied both linear biometry and geometric morphometrics and, thus, complemented previous studies that addressed the factors that affect variation in colouration (Saint-Girons et al., 1986; Martínez-Freiría et al., 2017) and in pholidotic (e.g. Bea et al., 1984; Martínez-Freiría and Brito, 2013) traits in this species. To our knowledge, this is the first study that implements geometric morphometric techniques to external head morphology in European vipers, providing a solid methodological toolkit for evaluating sources of head shape variation in these organisms (Fig. 3; Supp. Mat. 2). Interestingly, our analyses suggest a lack of sexual dimorphism, and lack of variation across colour morphs, in head shape. Instead, head shape variation is mainly driven by variation in size in this viper species. These results contrast with the patterns observed for linear biometric traits, for which we found important variation in size both between the sexes and across colour morphs. In some cases, variation in adult body proportions is determined by a marked differentiation in ontogenetic allometries, e.g. shaping tail length sexual dimorphism (Fig. 2). While both ontogenetic and static allometry analyses reflect the major contribution of size variation in shaping body relative dimensions, we also found that variation between the sexes and across colour morphs is still significant after size effects are taken into account, pointing to possible evolutionary mechanisms that may underlie intraspecific morphological variability in *Vipera seoanei*.

### Ontogeny and Sexual Dimorphism

Ontogenetic allometry is a biological phenomenon that occurs in different species, both endotherms and ectotherms, and can affect different aspects of organisms’ life (Herrel and Gibb, 2006; Wolf, Friggens and Salazar-Bravo, 2009). Our results clearly show that *Vipera seoanei* is no exception, as body size variation across ontogeny had an important contribution in shaping intraspecific morphological variability, both in body proportions (Table 2) and in head shape (Table 4). The detailed examination of ontogenetic allometric patterns performed here, provides a better understanding of the mechanisms that drive morphological variation in *Vipera seoanei*. For instance, we observed markedly different growth rates in the tail length of males and females. A decrease in growth rate after maturation has been reported for both sexes in other European vipers (e.g. *V. latastei*, Brito and Rebelo, 2003; *V. berus* in Bondarenko and Zinenko,2016). Such an effect is expected to be more accentuated in females, as considerable amount of the energy intake is dedicated to reproduction (Bonnet, Bradshaw and Shine, 1998; Bonnet et al., 2000). In our sample of *V. seoanei*, the tails of males, not only were longer compared to those of females, but also exhibited a faster growth rate (i.e. higher allometric slope) throughout all their ontogenetic trajectory. Having longer tails than females could be a benefit for male vipers, since other than containing the hemipenes, a longer tail could also be beneficial in increasing a male’s ability in courtship and/or male-male rivalry (King, 1989; Luiselli, 1996).

Examination of ontogenetic trajectories also provides a better understanding of head shape variation through geometric morphometric analyses. Indeed, we found that the supraocular region and the rear part of the head are the areas that undergo the highest variation during growth, where an increase in head size is accompanied by a reduction in relative eye size and an amplification of the posterior head region (Fig. 3). Presumably, these differences along the growth are concentrated on the non-ossified areas of the dorsal area of the head (Bellairs and Kamal 1981; Rieppel 1993; Polachowski and Werneburg 2013), which are less constrained and can be more intensely modified throughout ontogeny. Reinforcing this view, these are also the same areas that exhibit the largest portion of variation in shape across the whole sample (i.e. as revealed by Principal Components Analysis, Fig. 3), both before and after size effects are taken into account. Such variation, particularly that observed in the posterior area of the head, may be related to ontogenetic changes, as well as individual variation, in diet. As in other viper species (e.g. Pleguezuelos et al., 2007), juveniles of *V. seoanei* prey mainly on ectotherms of a relatively small size (e.g. lizards of the genera *Podarcis, Iberolacerta* and *Zootoca*), while adults mainly prey on endotherms (e.g. rodents of the genus *Microtus*; Braña, Bea and Saint Girons, 1988; Galán, 1988), which are larger. Being snakes gape-limited predators, the relative size of the head, as well as head proportions, may be correlated to relative prey size (Forsman and Shine, 1997; Bonnet et al., 2001; Lopez, Manzano and Prieto, 2013). Interestingly, such size-driven variation in global head shape occurs under a common allometric trend in both sexes (Table 4, Fig. 3), possibly pointing to shared ecological pressures in males and females of *V. seoanei*. Indeed, this pattern is in accordance with the lack of sexual differences in the diet of *V. seoanei* or other Iberian *Vipera* (e.g. Braña, Bea and Saint Girons, 1988; Martínez-Freiría et al., 2010).

Sexual dimorphism in head size and shape is a trait shared among many groups of reptiles and it has been usually associated with male antagonistic behaviour (Kaliontzopoulou, Carretero and Llorente, 2008; Barata et al., 2012; Kaliontzopoulou et al., 2012). In snakes, one of the possible hypotheses to explain head sexual dimorphism is related to male combats for access to females during the mating season (Tomović et al., 2002; Vincent, Herrel and Irschick, 2004; Andjelković, Tomović and Ivanović, 2016), which presumably occur in *V. seoanei*, as it was signaled for other *Vipera* species (Madsen, 1988; Senter, Harris and Kent, 2014). Accordingly, we expected to find more robust and larger heads in males than in females. Our results, however, showed that only snout height was significantly dimorphic once size effects were taken into account (Table 2, Fig. 2), while all the relative head dimensions were similar between males and females, as was the case for head shape. Further studies are needed to shed light into the functional role of snout height and its possible contribution in explaining the observed differences between the sexes.

### Colour Morphs

Differences in colouration and pholidotic traits have been suggested to be environmentally-driven in several viper species, including *V. seoanei* (e.g. Martínez-Freiría et al., 2010; Martínez-Freiría and Brito, 2013; Santos et al., 2014). Similarly, we hypothesized morphological differences across colour morphs could arise from distinct ecological pressures occurring in the main habitats occupied by them (i.e. lowland vs montane habitats). Indeed, the climatic conditions of lowland and montane habitats could lead to different durations of the annual cycle of vipers and, thus, distinct sizes and growth rates in individuals inhabiting these environments (e.g. Zuffi et al., 2009). Furthermore, distinct predation pressures at lowland and montane habitats (see Martínez-Freiría et al., 2017) might differentially affect survival, driving biometric differences among individuals from these environments. The patterns of variation observed for linear biometric traits of the four colour morphs of *V. seoanei* examined here may be partially explained according to this hypothesis.

Specifically, our results indicated that vipers of the *classic* morphotype were the smallest overall and did not exhibit sexual dimorphism in total body size. By contrast, females of *cantabrica* vipers, presented a higher SVL (Fig. 2, Supp. Mat. 3). Under the aforementioned environmental framework, a longer activity period in lowland habitats could allow *classic* females to reach sexual maturity sooner than females of *cantabrica*, being these females the smallest adults of all morphotypes. High predation pressure might be, however, eroding large-sized animals of *classic* (both sexes) in the lowlands, while low predation pressure might allow females of *cantabrica* to reach bigger sizes in the mountains. At this respect, *melanistic* vipers that also inhabit mountain environments, could profit from additional ecological advantages associated to a better thermal performance (e.g. increased growth rate, longer activity periods; Luiselli, 1992; Capula and Luiselli, 1994) and thus, show longer body sizes than the other colour morphs (Fig. 2). However, this result is not corroborated by our pairwise comparisons (Supp. Mat. 3). Interestingly, *bilineata* vipers, which occupy both coastal and montane habitats, show the largest variability for SVL, in our sample size likely as result of a combination of these ecological pressures occurring along a marked environmental gradient.

### Concluding remarks

Taken all together, our results provide a thorough description of morphological variability in *Vipera seoanei*, highlighting the relevance of combining different tools (i.e. linear and geometric morphometrics) and analyses that allow us to weight the contribution of different proximate and evolutionary factors in shaping intraspecific variation. As in most organisms (Klingenberg 2016), allometric variation and the effect of size on body proportions and shape constitute by far the most striking pattern observed in our data. By contrast, sexual differentiation – other than that in size – is limited to specific body parts, and in some cases emerges through sexually divergent growth trajectories (e.g. for tail length, Fig. 2), presumably related to sexual selection pressures acting on male individuals. Interestingly, our combined analyses also revealed the existence of complex patterns of biometric variation across colour morphs of *V. seoanei*, with potential relevance for ecological adaptation. Still, other factors not considered here could be involved in shaping morphological variability in *Vipera seoanei* and, thus, further studies addressing the relationship among morphological variability and environmental variables such as altitude, temperature or habitat are needed in order to explicitly test our hypotheses. Finally, although in our sample of *Vipera seoanei* head size was the predominant source determining head shape variation, without identified differences between the sexes or across colour morphs, our study puts forward a low-error analytical protocol for the implementation of geometric morphometric techniques in the investigation of head shape variation in European vipers.

## Acknowledgements

Friends and collaborators who helped in fieldwork. Stuff and curators from museum collections. This research is financed by the FEDER Funds through the COMPETE program and by Portuguese National Funds, within the scope of the project “PTDC/BIA-EVL/28090/2017-POCI-01-0145-FEDER-028090”. AK and FM-F are supported by contracts from FCT Portugal (IF/00641/2014/CP1256/CT0008 and DL57/2016/CP1440/CT0010, respectively).

## References

Adams, D.C., Collyer M.L., Kaliontzopoulou A. (2019): Geomorph: Software for geometric morphometric analyses. R package version 3.1.0. https://cran.r-project.org/package=geomorph.

Andjelković, M., Tomović, L., Ivanović, A. (2016): Variation in skull size and shape of two snake species (*Natrix natrix* and *Natrix tessellata*). Zoomorphology, 135: 243–253.

Arbour, J.H., Brown, C.M. (2014): Incomplete specimens in geometric morphometric analyses. Methods Ecol. Evol. 5: 16–26.

Badyaev, A.V. (2002): Growing apart: an ontogenetic perspective on the evolution of sexual size dimorphism. Trends Ecol. Evol. 17: 369–378.

Barata, M., Perera, A., Martínez-Freiría, F., Harris, D.J. (2012): Cryptic diversity within the Moroccan endemic day geckos *Quedenfeldtia* (Squamata: Gekkonidae): a multidisciplinary approach using genetic, morphological and ecological data. Biol. J. Linn. Soc. 106: 828–850.

Bea A., Bas S., Brañaa F., Saint-Girons H. (1984): Morphologie comparee et repartition de Vipera seoanei (Lataste, 1879), en Espagne. Amphibia-Reptilia 5:395–410.

Bellairs Ad’A, Kamal A.M. (1981): The chondrocranium and the development of the skull in recent reptiles. In: Gans C, Parsons TS, eds. Biology of the reptilia, Vol. 11: morphology F. London: Academic Press, 1–263.

Bondarenko, Z.S., Zinenko, O.I. (2016): Individual Growth Rates of Nikolsky’s Viper, Vipera berus nikolskii (Squamata, Viperidae) Vestnik Zoologii. Volume 50 (1): 65–70.

Bonduriansky, R. (2007): Sexual selection and allometry: a critical reappraisal of the evidence and ideas. Evolution (N. Y). 61: 838–849.

Bonduriansky, R., Day, T. (2003): The evolution of static allometry in sexually selected traits. Evolution 57: 2450–2458.

Bonnet, X., Bradshaw, S.D., Shine, R. (1998): Capital versus income breeding: an ectothermic perspective. Oikos 83: 333–341.

Bonnet, X., Naulleau, G., Shine, R., Lourdais, O. (2000): Reproductive versus ecological advantages to larger body size in female snakes, Vipera aspis. Oikos, 89(3), 509–518.

Bonnet, X., Shine, R., Naulleau, G., Thiburce, C. (2001): Plastic vipers: influence of food intake on the size and shape of Gaboon vipers (*Bitis gabonica*). J. Zool. 255: 341–351.

Bookstein, F.L. (1997): Landmark methods for forms without landmarks: morphometrics of group differences in outline shape. Med. Image Anal. 1: 225–243.

Braña, F. (2002): Vipera seoanei. In: Pleguezuelos JM, Marquez R, Lizana M (eds), Atlas y Libro Rojo de los Anfibios y Reptiles de Espana. Direccion General de Conservacion de la Naturaleza e Asociacion Herpetologica Espanola, Madrid, pp. 302–303.

Braña, F., Bea, A., Saint Girons, H. (1988): Composicion de la dieta y ciclos de alimentacion en *Vipera seoanei* Lataste, 1879. Variaciones en relacion con la edad y el ciclo reproductor. MUNIBE (Ciencias naturales), 40: 19–27.

Brito, J.C. (2008): *Vipera seoanei* Lataste, 1879. In: Loureiro A, Ferrand de Almeida N, Carretero MA, Paulo OS (eds), Atlas dos Anfíbios e Repteis de Portugal. Instituto da Conservacao da Natureza e da Biodiversidade, Lisboa, pp 184–185.

Brito, J.C., Rebelo, R. (2003): Differential Growth and Mortality Affect Sexual Size Dimorphism in *Vipera latastei*. Copeia 2003(4): 865–871.

Capula, M., Luiselli, L., 1994. Reproductive strategies in alpine adders, *Vipera berus*. The black females bear more often. Acta Oecol 15: 207–214.

Castella, B., Golay, J., Monney, J.C., Golay, P., Mebert, K., Dubey, S. (2013): Melanism, body condition and elevational distribution in the asp viper. J. Zool. 290: 273–280.

Claude, J. (2008): Morphometrics with R. Springer, New York.

Clusella-Trullas, S., Terblanche, J.S., Blackburn, T.M. and Chown, S.L. (2008): Testing the thermal melanism hypothesis: a macrophysiological approach. Funct. Ecol. 22: 232–238.

Clutton-Brock, T.H. (2007): Sexual selection in males and females. Science, 318: 1882–1885.

Collyer, M.L., Adams D.C. (2018): RRPP: An R package for fitting linear models to highdimensional data using residual randomization. Methods Ecol. Evol. 9: 1772–1779.

Collyer, M.L., Adams D.C. (2019): RRPP: Linear Model Evaluation with Randomized Residuals in a Permutation Procedure. https://cran.r-project.org/web/packages/RRPP.

Darwin, C. (1871): The descent of man and selection in relation to sex. London: J. Murray.

Dryden, I.L., Mardia, K.V. (1998): Statistical shape analysis. John Wiley & Sons, New York.

Fairbairn, D.J., Blanckenhorn, W.U., Szekely, T. (2007): Sex, size and gender roles. Evolutionary studies of sexual size dimorphism. New York: Oxford University Press.

Feldman, A., Meiri, S. (2013): Length–mass allometry in snakes. Biol. J. Linn. Soc. 108: 161–172.

Forsman, A. (1991): Adaptive variation in head size in *Vipera berus* L. populations. Biol. J. Linn. Soc. 43: 281–296.

Forsman, A. (1995a): Opposing fitness consequences of colour pattern in male and female snakes. J. Evol. Biol. 8: 53–70.

Forsman, A. (1995b): Heating rates and body temperature variation in melanistic and zigzag *Vipera berus*: Does colour make a difference? Ann. Zool. Fenn. 32: 365–374.

Forsman, A. (1996): An Experimental Test for food effects on head size allometry in juvenile snakes. Evolution 50: 2536–2542.

Forsman, A., Shine, R. (1997): Rejection of non-adaptive hypotheses for intraspecific variation in trophic morphology in gape-limited predators. Biol. J. Linn. Soc. 62: 209–223.

Gaffney, E.S. (1979): Comparative cranial morphology of recent and fossil turtles. Bull. Am. Mus. Nat. Hist. 164: 65–376.

Galán, P. (1988): Segregación ecológica en una comunidad de ofidios. Doñana Acta Vertebrata, 15: 59–78.

Gould, S.J. (1966): Allometry and size in ontogeny and phylogeny. Biol. Rev. 41: 587–640.

Greene, H. (1983): Dietary correlates of the origin and radiation of snakes. Am. Zool. 23: 431–441.

Gunz, P., Mitteroecker, P. (2013): Semilandmarks: A method for quantifying curves and surfaces. Hystrix 24: 103–109.

Gunz, P., Mitteroecker, P., Neubauer, S., Weber, G.W., Bookstein, F.L. (2009): Principles for the virtual reconstruction of hominin crania. J. Hum. Evol. 57: 48–62.

Harmon, L.J., Schulte II, J.A., Larson, A., Losos, J.B. (2003): Tempo and Mode of Evolutionary Radiation in Iguanian Lizards. Science 301: 961–964.

Henao-Duque, A.M., Ceballos, C.P. (2013): Sex-related head size and shape dimorphism in Mapaná snakes (*Bothrops asper*) kept in captivity. Rev. Colomb. Cienc. Pec. 26: 201–210.

Herrel, A., Gibb, A.C. (2006): Ontogeny of performance in vertebrates. Physiol. Biochem. Zool. 79: 1–6.

Kaliontzopoulou, A., Adams, D.C., van der Meijden, A., Perera, A., Carretero, M.A. (2012): Relationships between head morphology, bite performance and ecology in two species of Podarcis wall lizards. Evol. Ecol. 26: 825–845.

Kaliontzopoulou, A., Carretero, M.A., Llorente, G.A. (2008): Head shape allometry and proximate causes of head sexual dimorphism in *Podarcis* lizards: joining linear and geometric morphometrics. Biol. J. Linn. Soc. 93: 111–124.

Kaliontzopoulou, A., Pinho, C., Martínez-Freiría, F. (2018). Where does diversity come from? Linking geographical patterns of morphological, genetic, and environmental variation in wall lizards. BMC evolutionary biology, 18(1), 124.

King, R.B. (1989): Sexual dimorphism in snake tail length: sexual selection, natural selection, or morphological constraint? Biol. J. Linn. Soc. 38: 133–154.

King, R.B. (1997): Variation in brown snake (*Storeia dekayi*) morphology and scalation: sex, family, and microgeographic differences. J. Herpetol. 31: 335–346.

Klingenberg, C.P. (2016) Size, shape, and form: concepts of allometry in geometric morphometrics. Dev. Genes Evol. 226: 113–137.

Klingenberg, C.P., Barluenga, M., Meyer, A. (2002): Shape analysis of symmetric structures: quantifying variation among individuals and asymmetry. Evolution 56: 1909–1920.

Kodric-Brown, A., Sibly, R.M., Brown, J.H. (2006): The allometry of ornaments and weapons. PNAS 103: 8733–8738.

Krause, M., Burghardt, G., Gillingham, J. (2003): Body size plasticity and local variation of relative head and body size sexual dimorphism in garter snakes (*Thamnophis sirtalis*). J. Zool. 261: 399–407.

Lopez, M.S., Manzano, A.S., Prieto, Y.A. (2013): Ontogenetic Variation in Head Morphology and Diet in Two Snakes (Viperidae) from Northeastern Argentina. J. Herpetol. 47: 406–412.

Lovern, M.B., Holmes, M.M., Wade J. (2004): The Green Anole (*Anolis carolinensis*): A Reptilian Model for Laboratory Studies of Reproductive Morphology and Behavior, ILAR J. 45: 54–64.

Luiselli, L., (1992): Reproductive success in melanistic adders: a new hypothesis and some considerations on Andren and Nilson’s (1981) suggestions. Oikos 1992: 601–604.

Luiselli, L. (1996): Individual success in mating balls of the grass snake, *Natrix natrix*: size is important. J. Zool. 239: 731–740.

Madsen, T. (1988): Reproductive success, mortality and sexual size dimorphism in the adder, *Vipera berus*. Ecography 11: 77–80.

Martínez-Freiría, F., Brito, J.C. (2013): Integrating classical and spatial multivariate analyses for assessing morphological variability in the endemic Iberian viper *Vipera seoanei*. J. Zool. Syst. Evol. Res. 51: 122–131.

Martínez-Freiría, F., Brito, J.C. (2014): *Vipera seoanei* (Lataste, 1879). Fauna Ibérica. Vol. 10 Reptiles, 2a Edicion (eds M.A. Ramos), et al.), pp. 942-957. Museo Nacional de Ciencias Naturales - CSIC, Madrid.

Martínez-Freiría, F., De Lanuza, G.P.I., Pimenta, A.A., Pinto, T., Santos, X. (2017): Aposematism and crypsis are not enough to explain dorsal polymorphism in the Iberian adder. Acta Oecol. 85: 165–173.

Martínez-Freiría, F., Lizana, M., do Amaral, J.P., Brito, J. (2010): Spatial and temporal segregation allows coexistence in a hybrid zone among two Mediterranean vipers (Vipera aspis and V. latastei). Amphibia-Reptilia, 31 (2): 195–212.

Martínez-Freiría, F., Velo-Antón, G., Brito, J.C. (2015): Trapped by climate: interglacial refuge and recent population expansion in the endemic Iberian adder *Vipera seoanei*. Divers. Distrib. 21: 331–344.

McNamara, K.J. (2012): Heterochrony: The Evolution of Development. Evolution: Education and Outreach 5: 203–218.

Pleguezuelos, J., Brito, J., Fahd, S., Santos, X., Llorente, G., Parellada, X. (2007): Variation in the diet of the Lataste’s viper *Vipera latastei* in the Iberian Peninsula: seasonal, sexual and size-related effects. Anim. Biol. 57(1): 49–61.

Polachowski, K.M., Werneburg, I. (2013): Late embryos and bony skull development in *Bothropoides jararaca* (Serpentes, Viperidae). Zoology, 116 (1): 36–63.

R Development Core Team (2011) R: A language and environment for statistical computing. R Foundation for Statistical Computing, Vienna. http://www.R-project.org

Rieppel, O. (1993): Patterns of diversity in the reptilian skull. In the Skull, volume 2: Patterns of structural and systematic diversity. (J. Hanken and B. K. Hall, eds.). University of Chicago Press, Chicago, pp. 344–390.

Rodríguez, R.L., Cramer, J.D., Schmitt, C.A., Gaetano, T.J., Grobler, J.P., Freimer, N.B., Turner, T.R. (2015): The static allometry of sexual and nonsexual traits in vervet monkeys. Biol. J. Linn. Soc. 114: 527–537.

Rohlf, F.J. (1990): Morphometrics. Annu. Rev. Ecol. Syst. 21: 299–316.

Rohlf, F.J. (1999): Shape statistics: Procrustes superimpositions and tangent spaces. J. Class. 16: 197–223.

Rohlf, F.J. (2008): tpsDig version 2.29 Ecology and Evolution, SUNY at Stony Brook, NY.

Rohlf, F.J., Slice, D. (1990): Extensions of the Procrustes Method for the Optimal Superimposition of Landmarks. Syst. Zool. 39: 40.

Saint-Girons H., Braña F., Bea A. (1986): La distribucion de los diferentes fenotipos de Vipera seoanei Lataste, 1879, en la region de los Picos de Europa (Norte de la Península Iberica). Munibe 38: 121–128.

Santos, X., Azor, J. S., Cortés, S., Rodríguez, E., Larios, J., Pleguezuelos, J. M. (2017). Ecological significance of dorsal polymorphism in a Batesian mimic snake. Current zoology, 64(6), 745-753.

Santos, X., Vidal-García, M., Brito, J.C., Fahd, S., Llorente, G.A., Martínez-Freiría, F., Parellada, X., Plequezuelos, J.M., Sillero, N. (2014): Phylogeographic and environmental correlates support the cryptic function of the zigzag pattern in a European viper. Evol. Ecol. 28: 611–626.

Senter, P., Harris, S.M., Kent, D.L. (2014): Phylogeny of courtship and male-male combat behavior in snakes. PLoS ONE 9 (9): e107528.

Shine, R. (1989): Ecological causes for the evolution of sexual dimorphism: a review of the evidence. Q. Rev. Biol. 64: 419–461.

Shine, R. (1991): Intersexual dietary divergence and the evolution of sexual dimorphism in snakes. Am. Nat. 138: 103–122.

Shine, R. (1994): Sexual size dimorphism in snakes revisited. Copeia 2: 326–346.

Slatkin, M. (1984): Ecological Causes of Sexual Dimorphism. Evolution, 38: 622–630.

Tomović, L., Radojicic, J., Dzukic, G., Kalezic, M.L. (2002): Sexual dimorphism of the sand viper (*Vipera ammodytes* L.) from the central part of Balkan peninsula. Russ. J. Herpetol. 9: 69–76.

Valkonen, J.K., Niskanen, M., Björklund, M., Mappes, J. (2011): Disruption or aposematism? Significance of dorsal zigzag pattern of European vipers. Evol. Ecol. 25: 1047–1063.

Verwaijen, D., Van Damme, R., Herrel, A. (2002): Relationships between head size, bite force, prey handling efficiency and diet in two sympatric lacertid lizards. Funct. Ecol. 16: 842–850.

Vincent, S.E., Herrel, A., Irschick, D.J. (2004): Sexual dimorphism in head shape and diet in the cottonmouth snake (*Agkistrodon piscivorus*). J. Zool. 264: 53–59.

Wolf, M., Friggens, M., Salazar-Bravo, J. (2009): Does weather shape rodents? Climate related changes in morphology of two heteromyid species. Naturwissenschaften, 96: 93–101.

Zuffi, M. (2008): Colour pattern variation in populations of the European Whip snake, *Hierophis viridiflavus*: does geography explain everything? Amphibia-Reptilia, 29: 229–233.

Zuffi, M.A., Gentilli, A., Cecchinelli, E., Pupin, F., Bonnet, X., Filippi, E., Luiselli, L.M., Barbanera, F., Dini, F., Fasola, M. (2009): Geographic variation of body size and reproductive patterns in Continental versus Mediterranean asp vipers, *Vipera aspis*. Biol. J. Linn. Soc. 96 (2): 383–391.

