## supplemental material for "Sources of Intraspecific Morphological Variation in *Vipera seoanei*: Allometry, Sex, and Colour Phenotype"

*Amphibia-Reptilia*

CIBIO/InBIO, Centro de Investigação em Biodiversidade e Recursos Genéticos da Universidade do Porto. Instituto de Ciências Agrárias de Vairão. R. Padre Armando Quintas. 4485-661 Vairão Portugal;

### Supplementary Material

**Supplementary Material S1**

Table S1.1 - Number of individuals used for linear biometric analyses assigned to each group.

| SEX | AGE | bilineata | cantabrica | classic | melanistic | uniform | total |
|---|---|---|---|---|---|---|---|
| F | Juveniles | - | 3 | 5 | 3 | - | 11 |
| | Subadults | - | 4 | 2 | 1 | - | 7 |
| | Adults | 15 | 56 | 92 | 23 | 5 | 173 |
| M | Juveniles | 1 | 15 | 15 | 1 | - | 32 |
| | Subadults | 2 | 14 | 20 | 4 | - | 40 |
| | Adults | 13 | 63 | 102 | 21 | 1 | 128 |

Table S1.2 - Number of individuals used for geometric morphometric analyses assigned to each group.

| SEX | AGE | bilineata | cantabrica | classic | melanistic | uniform | total |
|---|---|---|---|---|---|---|---|
| F | Juveniles | - | 1 | 3 | - | - | 4 |
| | Subadults | - | 1 | 1 | - | - | 2 |
| | Adults | 4 | 20 | 25 | 3 | - | 52 |
| M | Juveniles | - | 8 | 6 | - | - | 14 |
| | Subadults | - | 3 | 8 | - | - | 11 |
| | Adults | 3 | 15 | 19 | 4 | - | 41 |

Table S1.3 - Total and by group number of missing values replaced for each trait: SVL (snout-vent length), TAIL (tail length), HL (head length), ML (mouth length), JL (jaw length), HW (head width), HH (head height), SH (snout height), TOTAL (total sample size).

| | Total | Juveniles | | Subadults | | Adults | |
|---|---|---|---|---|---|---|---|
| | | M | F | M | F | M | F |
| SVL | - | - | - | - | - | - | - |
| TAIL | 11 | - | - | 1 | - | 4 | 6 |
| HL | 5 | 2 | - | 2 | - | 1 | - |
| ML | 20 | 4 | - | 4 | - | 7 | 5 |
| JL | 24 | 4 | 1 | 3 | - | 7 | 9 |
| HW | 55 | 2 | 1 | 7 | - | 23 | 22 |
| HH | 73 | 2 | 2 | 12 | 1 | 31 | 25 |
| SH | 50 | 2 | 2 | 8 | 1 | 21 | 16 |
| TOTAL | 391 | 32 | 11 | 40 | 7 | 128 | 173 |

Table S1.4 - Minimum (Min), mean, maximum (Max) and standard deviation (SD) values for linear morphometric traits: SVL (snout-vent length), TAIL (tail length), HL (head length), ML (mouth length), JL (jaw length), HW (head width), HH (head height), SH (snout height).

| | M | | | | F | | | |
|---|---|---|---|---|---|---|---|---|
| | Min | Mean | Max | Sd | Min | Mean | Max | Sd |
| **SVL** | 145 | 342.876 | 490 | 86.274 | 145 | 396.068 | 535 | 64.752 |
| **TAIL** | 20 | 54.842 | 85 | 14.774 | 21 | 51.117 | 79 | 8.710 |
| **HL** | 9.060 | 15.535 | 20.790 | 2.467 | 9.350 | 16.872 | 20.520 | 2.001 |
| **ML** | 9.730 | 16.343 | 21.840 | 2.668 | 10.160 | 17.760 | 22.400 | 1.947 |
| **JL** | 10.920 | 19.088 | 25.640 | 3.215 | 11.620 | 20.918 | 25.500 | 2.328 |

| | | | | | | | | |
|---|---|---|---|---|---|---|---|---|
| **HW** | 5.500 | 12.386 | 20.620 | 2.216 | 6.560 | 13.625 | 18.880 | 1.918 |
| **HH** | 4.200 | 7.419 | 10.350 | 1.274 | 4 | 8.241 | 13.050 | 1.152 |
| **SH** | 1.360 | 2.856 | 3.980 | 0.492 | 1.740 | 3.043 | 4.300 | 0.377 |

**Supplementary Material S2**

Evaluation of measurement error during digitizing landmarks to study variation in head shape in *Vipera seoanei*.

*Methods*

In order to examine the effect of digitizing error in studying intraspecific head shape variation in *Vipera seoanei*, one of us (GA) digitized all specimens twice. To test for the presence of measurement error and evaluate its magnitude as compared to the total shape variation present in our sample, we first superimposed all original and repeated landmark configurations. and then we fit a linear model on Procrustes residuals with individual (ind) and digitizing repetition (rep) as explanatory variables. Linear models were fitted using the function *procD.lm* of *geomorph* R-package (Adams, Collyer and Kaliontzopoulou, 2019) and evaluated using residual randomization procedures as implemented in the R-package *RRPP* (Collyer and Adams 2018, 2019).

To further verify the consistency of both digitizations, we used a two-block partial least-squares analysis (PLS: Rohlf and Corti 2000), to quantify the multivariate association between shape variables produced in each of them. The significance of the correlation coefficient between PLS vectors was evaluated through 1000 data permutations. All PLS procedures were implemented using the function *two.b.pls* of *geomorph* R-package (Adams, Collyer and Kaliontzopoulou, 2019).

Finally, to investigate whether specimen origin (museum collection vs. alive vs. road killed) had an effect on the precision of shape data collection, we first quantified individual measurement error (indME) as the Procrustes distance between the two digitizations available for each specimen. Using these individual values, we then fit a linear model with indME as the response variable and specimen origin as the predictor, the significance of which was also evaluated using RRPP as described above.

*Results*

ANOVA statistics derived from the linear model fit to evaluate digitizing error indicated the occurrence of significant shape variation both among individuals and between the two digitization rounds (Table S1). Examination of $R^2$ values related to different effects suggests that 91.5% of shape variation is attributable to differences among individuals, while the rest is due to the effect of measurement error. Of this, about 3.4% is systematic error (repetition effect). By contrast, about 5.1% is random error, where digitization effects vary among individuals, as indicated by the residual term, which in this case corresponds to the ind×rep interaction. This means that systematic error, which may bias downstream shape analysis, represents – in our case – less than 5% of the total shape variation in our sample.

**Table S2.1:** ANOVA statistics from a linear model fit to shape data to evaluate the effect of individual (ind) and digitizing repetition (rep) on shape variation. df: degrees of freedom, SS: Sums of Squares, MS: Mean Squares, $R^2$: coefficient of determination, F: F-value, Z: standardized z-score, p: corresponding p-value based on 1000 permutations of residuals.

| | **df** | **SS** | **MS** | **$R^2$** | **F** | **Z** | **p** |
|---|---|---|---|---|---|---|---|
| **ind** | 139 | 1.439 | 0.010 | 0.915 | 17.958 | 22.370 | 0.001 |
| **rep** | 1 | 0.054 | 0.054 | 0.034 | 93.463 | 6.426 | 0.001 |
| **Residuals** | 139 | 0.080 | $5.77*10^{-4}$ | 0.051 | | | |
| **Total** | 279 | 1.573 | | | | | |

The partial least-squares analysis implemented to examine the multivariate association between the shape variables produced by both digitizing repetitions confirms this observation. Indeed, the two-blocks PLS provided a significant vector of association between shape variable blocks, where the correlation among vectors was $r_{PLS} = 0.939$ ($p = 0.001$).

Finally, our test of whether individual measurement error depended on the origin of specimen collection, including 98 specimens from museum collections, 35 collected alive in the field, and 8 specimens found as road kills, indicated no significant effect (Table S2.2).

**Table S2.2:** ANOVA statistics from a linear model fit to individual measurement error (indME) to evaluate the effect of specimen origin (ori). df: degrees of freedom, SS: Sums of Squares, MS: Mean Squares, $R^2$: coefficient of determination, F: F-value, Z: standardized z-score, p: corresponding p-value based on 1000 permutations of residuals.

| | **Df** | **SS** | **MS** | **$R^2$** | **F** | **Z** | **p** |
|---|---|---|---|---|---|---|---|
| **ori** | 2 | 0.000 | 0.000 | 0.010 | 0.713 | 0.182 | 0.477 |
| **Residuals** | 137 | 0.033 | 0.000 | 0.990 | | | |
| **Total** | 139 | 0.033 | | | | | |

Given these results, we consider that, despite the existence of digitizing error, which is practically unavoidable in GM studies (Arnqvist and Mårtensson, 1998), the magnitude of systematic error is very reduced as compared to the total shape variation present in our data, and that the origin of the specimens is unlikely to influence substantially the shape patterns observed here.

*References*

Arnqvist. G., Mårtensson. T. (1998): Measurement error in geometric morphometrics: Empirical strategies to assess and reduce its impact on measures of shape. Acta Zool. Acad. Sci. Hungaricae **44**: 73-96.

Adams. D.C., Collyer M.L., Kaliontzopoulou A. (2019): Geomorph: Software for geometric morphometric analyses. R package version 3.1.0. https://cran.r-project.org/package=geomorph.

Collyer M.L., Adams D.C. (2018): RRPP: RRPP: An R package for fitting linear models to high-dimensional data using residual randomization. Methods in Ecology and Evolution. **9**(2): 1772-1779.

Collyer M.L., Adams D.C. (2019): RRPP: Linear Model Evaluation with Randomized Residuals in a Permutation Procedure. https://cran.r-project.org/web/packages/RRPP.

Rohlf. F.J., Corti M. (2000): The use of partial least-squares to study covariation in shape. Syst. Biol. **49**: 740-753.

**Supplementary Material S3**

Post-hoc comparisons for morphological variables on which ANOVA comparisons indicated a significant effect of colour morph (Table 2). Pairwise comparisons based on 1000 random permutations of Euclidean distances were performed across pattern × sex groups, when both variables had a significant effect (i.e. SVL, TAIL, ML); or between pattern groups only, when the effect of sex was not significant (i.e. HL). Significant pairwise differences (at a = 0.05) are highlighted in boldface.

| **SVL** | bilineata F | cantabrica F | classic F | melanistic F | bilineata M | cantabrica M | classic M | melanistic M |
|---|---|---|---|---|---|---|---|---|
| bilineata F | | | | | | | | |
| cantabrica F | 0.571 | | | | | | | |
| classic F | 0.922 | **0.043** | | | | | | |
| melanistic F | 0.448 | 0.368 | 0.068 | | | | | |
| bilineata M | 0.157 | 0.120 | 0.370 | 0.093 | | | | |
| cantabrica M | 0.171 | 0.060 | 0.444 | 0.066 | 0.405 | | | |
| classic M | 0.629 | 0.639 | 0.990 | 0.484 | 0.077 | 0.199 | | |
| melanistic M | 0.783 | 0.095 | 0.722 | 0.091 | 0.479 | 0.588 | 0.983 | |

| **TAIL** | bilineata F | cantabrica F | classic F | melanistic F | bilineata M | cantabrica M | classic M | melanistic M |
|---|---|---|---|---|---|---|---|---|
| bilineata F | | | | | | | | |
| cantabrica F | 0.532 | | | | | | | |
| classic F | 0.127 | 0.2795 | | | | | | |
| melanistic F | 0.468 | 0.758 | 0.553 | | | | | |
| bilineata M | 0.446 | 0.724 | 0.776 | 0.876 | | | | |
| cantabrica M | 0.456 | 0.230 | 0.051 | 0.290 | 0.383 | | | |
| classic M | 0.345 | 0.186 | **0.013** | 0.242 | 0.347 | 0.958 | | |
| melanistic M | 0.435 | 0.652 | 0.827 | 0.827 | 0.91 | 0.374 | 0.339 | |

| **ML** | bilineata F | cantabrica F | classic F | melanistic F | bilineata M | cantabrica M | classic M | melanistic M |
|---|---|---|---|---|---|---|---|---|
| bilineata F | | | | | | | | |
| cantabrica F | 0.255 | | | | | | | |
| classic F | 0.368 | 0.274 | | | | | | |
| melanistic F | 0.185 | 0.102 | 0.956 | | | | | |
| bilineata M | 0.281 | 0.182 | 0.610 | 0.637 | | | | |
| cantabrica M | 0.447 | 0.632 | 0.279 | 0.103 | 0.211 | | | |
| classic M | 0.297 | 0.186 | 0.606 | 0.626 | 0.929 | 0.216 | | |
| melanistic M | 0.886 | **0.029** | 0.797 | 0.674 | 0.818 | 0.554 | 0.895 | |
